# Mechanisms of cortical microtubule organization in epidermal keratinocytes

**DOI:** 10.1101/2024.10.08.617151

**Authors:** Keying Guo, Andreas Merdes

## Abstract

Microtubules in many differentiated cell types are reorganized from a radial, centrosome-bound array into a cell type-specific, non-centrosomal network. In epidermal keratinocytes, cortical microtubules are organized from desmosomes (Lechler, T., and Fuchs, E. 2007. J. Cell Biol. 176:147-154). Details of this organization are poorly understood. We used immunofluorescence expansion microscopy to visualize directly the contact between cortical microtubules and desmosomes in murine skin tissue. Microtubule bound laterally to desmosomes, or with their ends at mixed polarity. Experiments including time-lapse microscopy of EB3-GFP, microtubule regrowth after depolymerization, and expression of ectopic ninein that was sequestered to the plasma membrane with a CAAX sequence motif, indicated that limited nucleation of microtubules can occur at the cortex, but mostly, cortical microtubules may accumulate by translocation from non-cortical sites and anchorage to desmosomes.

## Introduction

At the interface between the organism and the environment, the skin provides mechanical protection and constitutes an important barrier that prevents the invasion of pathogens, the unintentional entrance of exogenous substances such as allergens or toxins, and the uncontrolled loss of body water and solutes. The barrier function of the skin is mainly attributed to the epidermis, a stratified epithelium, with a basal layer of proliferative cells and three suprabasal compartments of differentiated keratinocytes: spinous, granular and cornified layers. Differentiated keratinocytes in the spinous layer assemble massive amounts of cytoskeletal filaments, in particular intermediate filaments of the keratins 1 and 10 (Fuchs, 1990). At the same time, keratinocytes form numerous intercellular junctions, comprising desmosomes that act as anchorage points for the keratin fibers, as well as tight junctions and adherens junctions (Sumigray & Lechler, 2015). Together, the network of keratins and intercellular junctions provides stability and optimal elasticity of the epithelium and represents a mechanical barrier. In addition, keratinocytes in the granular layer form a lipid-rich hydrophobic shield by secreting small membranous organelles, the lamellar bodies. The secretion of lamellar bodies, as well as the assembly and maintenance of intercellular junctions depend in part on microtubule-dependent transport: lamellar bodies interact with the microtubule-binding protein CLIP-170 (Raymond et al., 2008), and secretion of lamellar bodies necessitates the presence of the dynein-activator ninein (Redwine et al., 2017; Lecland et al., 2019). Moreover, microtubules are involved in the delivery of junctional proteins to the assembly sites of tight junctions, adherens junctions, and desmosomes (Vasileva & Citi, 2018). It has been proposed that the delivery of the desmosomal proteins desmoglein 2 and desmocollin 2 occurs along microtubules that are anchored at the cell cortex (Nekrasova et al., 2011). The concept of cortical microtubule anchorage in keratinocytes is further supported by the discovery of a microtubule-organizing role of the desmosome protein desmoplakin (Lechler & Fuchs, 2007). Desmoplakin is thought to bind microtubules indirectly, by recruiting proteins that are localized at the centrosome prior to keratinocyte differentiation, including ninein, Lis1, and Ndel1 (Lechler & Fuchs, 2007; Sumigray et al., 2011). This reorganization of microtubules is believed to be essential for the functionality of the epidermal barrier (Sumigray et al., 2011). Consistently, keratinocytes fail to accumulate microtubules at the cell cortex in the absence of ninein, and the skin of ninein-knockout mice displays barrier defects during development (Lecland et al., 2019).

So far, the mechanisms of cortical microtubule assembly and the principles of cortical microtubule organization in keratinocytes are only poorly understood. Whereas numerous experiments were performed in cell cultures, only few studies were carried out in skin tissue (Lechler & Fuchs, 2007; Sumigray et al., 2011; 2012; Muroyama & Lechler, 2017). Moreover, the origin of cortical microtubules remains unknown: it is unclear whether microtubules are nucleated directly at the cortex or whether they originate from non-cortical sites, following capture and/or stabilization at cortical sites. In this manuscript, we present immunofluorescence of microtubules and microtubule-end markers in mouse epidermis and in mouse keratinocyte cultures, combined with expansion microscopy and immuno-electron microscopy, to visualize directly the interaction of microtubules with the keratinocyte cortex at high resolution. Furthermore, we provide evidence for multiple origins of these microtubules, cortical and non-cortical, based on depolymerization and re-growth experiments, and transfection experiments with cortically tagged ninein.

## Materials and methods

### Cell culture

NIH-3T3 cells were cultured in DMEM containing 10% fetal bovine serum. MPEK-BL6 cells (CELLnTEC) were cultured in CnT-PR medium (CELLnTEC) for proliferation. Differentiation was induced near confluence, by replacing the medium with CnT-PR-D (CELLnTEC), and by adding 1.2mM CaCl_2_ for two days of culture.

### Expansion microscopy of mouse skin

Samples were collected from the tails of newborn C57Bl/6 mice. Samples were fixed for 5 hours in 4% paraformaldehyde in 60mM PIPES, 25 mM HEPES, 1 mM EGTA, 2 mM MgCl_2_ (PHEM), pH 6.9, at room temperature, rinsed in buffer, transferred into Tissue-Tek O.C.T. compound (Sakura), and subsequently frozen. Cryosections of 10 mm thickness were attached to glass coverslips and prepared for immunofluorescence. If formaldehyde fixation masked the antigens, samples were directly embedded in O.C.T. compound. Expansion microscopy was performed using a modified protocol from Gambarotto et al. (2019). Briefly, cryosections on coverslips were post-fixed in a mixture of 0.7% formaldehyde and 1% acrylamide in PBS for 4 hours at 37°C, followed by embedding in 10% acrylamide, 0.1% bis-acrylamide, 23% acrylic acid in PBS. Polymerization of the embedding mixture was started by the addition of 0.5% ammonium peroxodisulfate and 0.5% tetramethylethylenediamine on ice, and after 5 minutes followed by transfer to 37°C for 1 hour. Fully polymerized samples were treated for 90 minutes at 95°C with a denaturing buffer containing 50mM Tris/HCl, 0.2M NaCl, 5.7% sodium dodecyl sulfate, pH 9.0. Expansion of samples was performed by repeated soaking in fresh 100 ml volumes of deionized water for 1 hour. The samples were reproducibly found to expand by a factor of 4.1. For immunofluorescence, samples were incubated for 30 minutes in PBS, followed by standard immunofluorescence protocols, but with 4x concentrated antibody solutions, as compared to regular, non-expansion protocols. Following antibody staining, samples were post-fixed once more in 4% formaldehyde in PBS, and final expansion was achieved in deionized water.

Samples were analyzed on a Zeiss Axiovert 200M microscope using 100x/1.4NA, 63x/1.4NA, and 40x/1.3NA objectives, a Zeiss AxioCam MRm camera, and Zeiss Axiovision software. Stacks of images at 0.2-0.3 mm distance were acquired with the help of a Z-motor. Deconvolution of image stacks was performed with Huygens Professional software (Scientific Volume Imaging). Unless indicated otherwise, single optical sections were selected for the figures of this manuscript. The shift between images at different wavelengths, due to deviations of filter cube alignments and chromatic aberration, was quantified in supplemental Figure 1A.

### Immunoelectron microscopy

Differentiated MPEK cells on glass coverslips were fixed for 10 minutes at room temperature in 0.5% glutaraldehyde in PHEM, pH 6.9. Following extraction with 0.5% Triton-X-100 in PHEM for 15 minutes, cells were rinsed with buffer and treated twice for 15 minutes with freshly prepared 0.1% sodium borohydride in PBS at room temperature, to reduce reactive aldehyde groups. After rinsing with buffer, coverslips were incubated for 30 minutes at room temperature in blocking buffer, containing 0.8% bovine serum albumin and 0.1% fish skin gelatin in PBS. Cells were stained with primary antibody DM1A (Sigma-Aldrich), diluted 1:200 in blocking buffer, for two hours in a humidified chamber at room temperature. After rinsing three times 10 minutes with PBS, cells were incubated with rabbit anti-mouse IgG (Rockland) at 2mg/ml in blocking buffer, for two hours at room temperature. Following three more rinses with PBS, cells were incubated overnight at room temperature with protein A coupled to 5nm gold (CMC ImmunoGold Reagents, UMC Utrecht, NL), diluted 1:50 in blocking buffer. After three more rinses with buffer, cells were post-fixed for 5 minutes with 1% glutaraldehyde in PBS, rinsed in deionized water, and submitted to a series of dehydration in mounting concentrations of ethanol. Following infiltration for 30 minutes with a 1:1 mixture of araldite CY212 (Agar Scientific) in dehydrated ethanol, and three times 1 hour with pure araldite, coverslips were flat-embedded in araldite on glass slides, using small coverslip fragments as spacers, for 16 hours at 60°C. Slices of cells in polymerized araldite were recovered with a scalpel, after removal of the coverslip surface with 40% hydrofluoric acid and excessive rinses with deionized water. These slices were mounted on araldite blocks with cyanoacrylate glue, followed by ultrathin sectioning, and staining on copper grids with 5% uranyl acetate for 10 minutes, and lead citrate (Reynolds, 1963), for 5 minutes.

### Antibodies

The following antibodies were used for immunofluorescence microscopy: mouse-anti-alpha-tubulin DM1A (Sigma-Aldrich), 1:1000; guinea pig anti-alpha and anti-beta-tubulin monobodies AA345 and AA344 (ABCD antibodies scFv-F2C & scFv-S11B) 1:250; mouse anti-gamma-tubulin Tu-30 (Exbio), 1:200; rabbit anti-gamma-tubulin R75 (Julian et al., 1993), 1:200; mouse anti-GCP6 H9 (Santa Cruz Biotechnology), 1:60; mouse anti-EB1 (BD Biosciences), 1:250; rabbit anti-Cep170 (Abcam), 1:100; rabbit anti-CAMSAP2 #17880-1-AP (Proteintech), 1:300; rabbit anti-CLASP1 (Proteintech), 1:100; rabbit anti-desmoplakin #25318-1-AP (Proteintech), 1:100; mouse anti-desmoplakin 11-5F (Sigma-Aldrich), 1:20; mouse anti-dynein intermediate chain, clone 70.1 (Sigma-Aldrich), 1:500; rabbit anti-GM130 (Abcam), 1:500; rabbit anti-Lis1 (Santa Cruz Biotechnology), 1:200; mouse anti p150 Glued dynactin (BD Biosciences), 1:250; rabbit anti-Cdk5rap2 (Srsen et al., 2009), 1:100; rabbit anti-human ninein, raised against amino acids 1709-2090 (Uniprot Q8N4C6-1; Srsen et al., 2009), 1:80. A new rabbit serum was raised against mouse ninein, by cloning cDNA encoding mouse ninein (Uniprot Q61043-3, amino acids 1-496) into vector pRSET-A (Invitrogen), to produce a fusion protein with an amino-terminal hexa-histidine-tag. The protein was purified over Ni-NTA-Agarose (Qiagen), concentrated and dialyzed against PBS, prior to immunization (Kaneka Eurogentech). For immunofluorescence, the serum was diluted 1:80. Characterization of the serum is documented in supplemental Figure 1B.

### Microtubule regrowth assay

To depolymerize microtubules in MPEK-BL6 cells, culture medium was supplemented with 3.3 mM nocodazole, and cultures were incubated on ice for 3 hours. Nocodazole was then washed off twice for 30 minutes with fresh medium at 0°C, and microtubule regrowth was induced by transferring cultures to 37°C.

### Transfection experiments

NIH-3T3 cells were transiently transfected using Lipofectamine 2000 (Invitrogen), according to the manufacturer’s instructions. The following plasmids were used: a pEGFP-ninein plasmid (Delgehyr et al., 2005) was modified by inserting nucleotides encoding the terminal 14 amino acids of human KRas in front of the stop codon (EGFP-ninein-CAAX). In an alternative approach, an anti-GFP nanobody (Caussinus et al., 2011) was tagged with the CAAX motif from KRas, and the nanobody-encoding plasmid was co-transfected with ninein, tagged with EGFP at its carboxy-terminus. In separate experiments, spastin constructs pEYFP-mSpastin C530 and pEYFP-mSpastin C530 mut1A (Lacroix et al., 2010) were co-transfected with EGFP-ninein-CAAX. These cells were fixed 6 hours after transfection in methanol at -20°C, stained for immunofluorescence of alpha-tubulin, and identified by fluorescence microscopy of the EGFP and EYFP tags. EYFP signal was detected using a filter set with a 510 nm excitation filter, a Triband dichroic mirror, and a 542 nm emission filter. MPEK cells were transfected after 6 days of differentiation with EB3-GFP (Mogessie et al., 2015) and followed by time-lapse microscopy at 10s intervals.

## Results

### Lateral and end-on contacts of microtubules at the keratinocyte cortex

To visualize microtubule organization in skin tissue, we prepared cryosections of skin samples from the tails of newborn mice. Conventional immunofluorescence produced blurred images of the microtubules (Fig. 1A). To obtain improved images, we applied a protocol of expansion microscopy (Gambarotto et al., 2019) that expanded our samples isometrically by a factor of 4.1 and that allowed the resolution of individual microtubules and desmosomes. Details of the desmosomal structure were visualized, including the two cytoplasmic plaques, interspaced by the extracellular desmoglea (Fig. 1B). Cortical microtubules were seen in the proximity of the cytoplasmic plaques, touching these either end-on, or running parallel to the plaques within a distance of approximately 100 nm (Fig. 1C).

**Figure 1:**
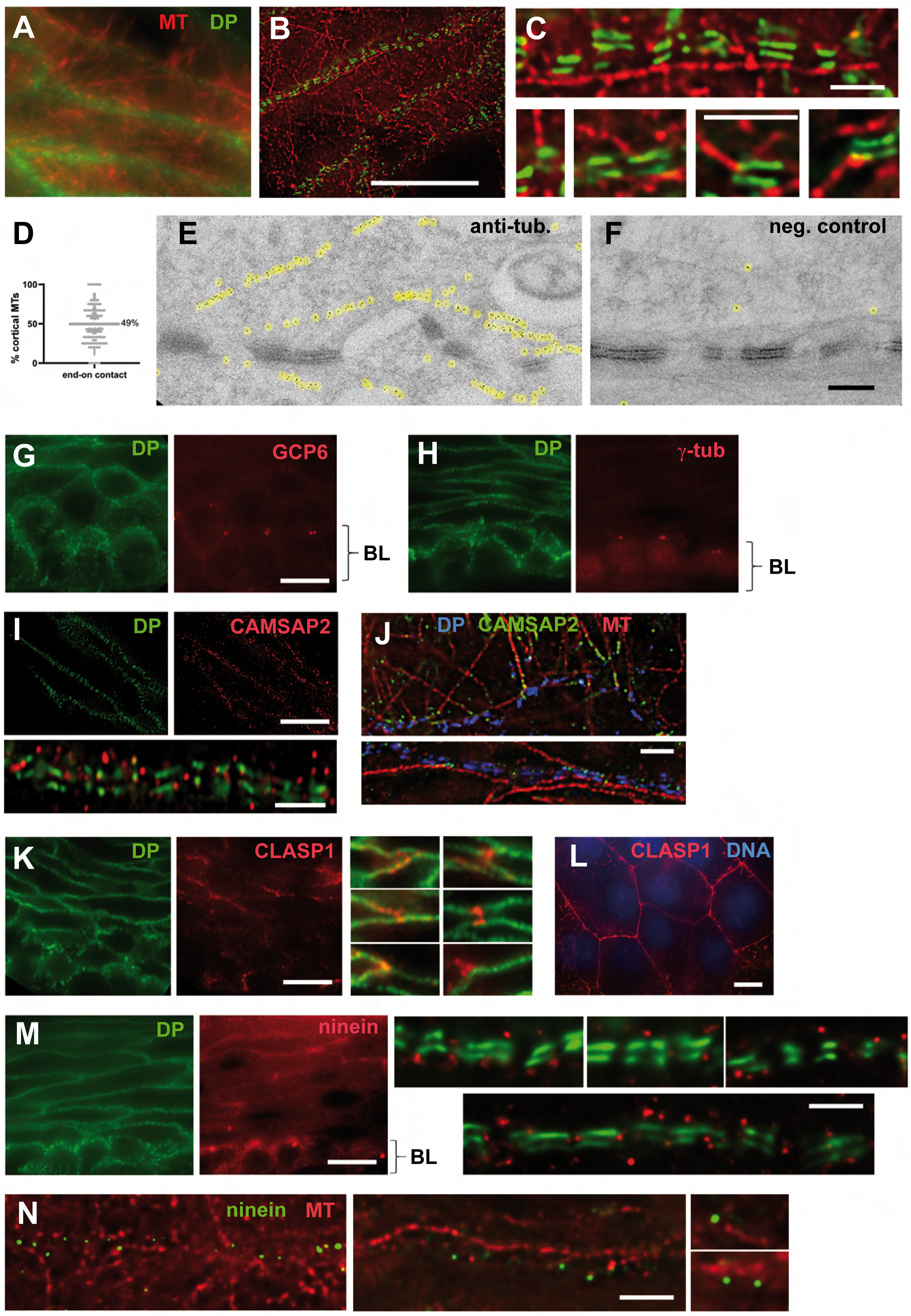
Cortical microtubule organization in epidermal keratinocytes and in differentiated keratinocyte cultures. (A) Cryosection of the epidermis from newborn mouse tail, stained with monobodies against alpha-and beta-tubulin (MT, red) and desmoplakin (DP, green). Cells from the spinous layer are shown, bordering the granular layer. (B) Cryosection as in (A), but subjected to an expansion microscopy protocol. (C) Higher magnification of expansion microscopy samples, stained as in (A). A microtubule running parallel to the cortex is shown (top), as well as several microtubules with end-on contacts to desmosomes (bottom). (D) Percentage of microtubules with end-on contacts to desmosomes. Microtubules were analyzed in cortical segments of micrographs from epidermis. Each segment had a length of 26 mm after expansion, displaying on average 5 microtubules. The percentage of microtubules with either end-on contact or lateral contact was determined for each segment. N=311 segments were analyzed, each data point corresponding to one segment. (E) Immuno-electron microscopy of alpha-tubulin in differentiated MPEK cells. The microtubules were detected with 5nm immuno-gold, and each gold particle was highlighted by a yellow background, using the color balance tool of Adobe Photoshop. (F) Control section without primary antibody, underlining the specificity of alpha-tubulin/immuno-gold staining in (E). (G) Cryosection of the epidermis from newborn mouse tail, stained with antibodies against desmoplakin and the gamma-TuRC component GCP6. The location of the basal layer (BL) is indicated. GCP6 localizes to the centrosomes in the basal layer. (H) Cryosection as in (G), stained with antibodies against desmoplakin and gamma-tubulin (g-tub). Gamma-tubulin localizes to the centrosomes in the basal layer. (I) Cryosection of epidermis after expansion, stained with antibodies against desmoplakin (green) and CAMSAP2 (red). Bottom, enlarged view of a cortical segment of expanded epidermis. (J) Cortical areas of differentiated MPEK cells after expansion, stained with antibodies against desmoplakin (DP, blue), CAMSAP2 (green), and alpha-tubulin (MT, red). (K) Cryosection of the epidermis from newborn mouse tail, stained with antibodies against desmoplakin (green) and CLASP1 (red). On the right, enlarged views of tricellular and tetracellular contacts in the epidermis are shown. (L) Differentiated MPEK cells, stained for CLASP1 (red) and DNA (blue). (M) Cryosection of the epidermis, stained with antibodies against desmoplakin (green), and ninein (red). The basal layer (BL) is indicated. Ninein staining at the cortex is most pronounced from the 3^rd^ suprabasal layer upwards. On the right, enlarged segments of the epidermal keratinocyte cortex are shown after expansion. (N) Cortical regions of keratinocytes in epidermal cryosections after expansion, stained with antibodies against ninein (green) and alpha-tubulin (red). Bars, (B, G, I, K, L, M) 10mm, (C, J, N, and enlarged areas in I, M) 1mm, (F) 0.2mm. The bar sizes in expanded samples correspond to the regular dimensions before expansion, i.e. the measured dimensions in the micrographs were divided by the expansion factor of 4.1.

We quantified the relative abundance of the two microtubule populations on 311 segments of the plasma membrane of spinous layer keratinocytes, and found nearly equal percentages (49% of end-on contacts, 51% of lateral contacts; see graph in Fig. 1D). To verify microtubule organization by an independent method, we employed immuno-electron microscopy, using antibody against tubulin, detected by protein A-immunogold. First attempts on fixed skin tissue failed, due to the very limited permeability of the material for immunogold (unpublished observations). Conventional transmission electron microscopy was equally complicated by the fact that the high abundance of keratin fibers in the tissue obscured the less abundant microtubules (unpublished observations). We finally succeeded by investigating cortical areas of MPEK keratinocyte cultures after differentiation *in vitro*, confirming the existence of cortical microtubules in the immediate proximity to desmosomes (Fig. 1E, F), and validating the data from expansion microscopy, since similar dimensions for desmosomes and microtubule distances were found.

The detection of microtubule-ends at the keratinocyte cortex immediately raised the question of their origin and of their polarity. To test whether microtubule nucleation takes place at or near desmosomes, we performed immunofluorescence of the gamma-tubulin ring complex, using antibodies against GCP6 and gamma-tubulin. Although both proteins displayed centrosomal localization in cells of the basal layer, this staining was largely lost in keratinocytes in the spinous layer, and no specific staining was detected at the keratinocyte cortex (Fig. 1G, H). Because earlier observations have indicated that centrosomal microtubule anchorage is lost in differentiated keratinocytes (Lechler & Fuchs, 2007; Muroyama et al., 2016), we tested whether the microtubule-ends next to desmosomes represent non-centrosomal minus-ends of microtubules protected by proteins of the patronin/CAMSAP family (Toya & Takeichi, 2016). We confirmed that CAMSAP2 is indeed enriched at the keratinocyte cortex near desmosomes in skin (Fig. 1I). We further observed that part of the microtubule-ends that attached to cortical sites at or near desmosomes in MPEK cells were decorated by comets of CAMSAP2 (Fig. 1J). However, not all cortical microtubule-ends were CAMSAP2-positive, and microtubules running parallel to the cortex showed no CAMSAP2 enrichment along their lattice. For this reason, we wanted to test whether microtubule-plus-ends were equally present at the cortex of skin keratinocytes. Immunofluorescence with antibodies against the plus-end marker EB1 failed to produce any staining in skin tissue, likely due to epitope masking after aldehyde fixation (unpublished observation). Likewise, immunofluorescence of skin sections embedded in O.C.T. compound without fixation failed to provide any EB1 signal, likely due to loss of dynamic microtubules during the embedding and freezing process of the tissue (unpublished observation). We obtained indirect information on microtubule orientation by investigating the presence of plus-end-stabilizing CLASP proteins that can bind to the cortex independent of microtubules (Mimori-Kiyosue et al., 2005; Lansbergen et al., 2006). A CLASP1 antibody revealed uneven localization at the cortical region of suprabasal keratinocytes (Fig. 1K). Closer inspection suggested that CLASP1 is particularly enriched near tricellular junctions. In MPEK cells, on the other hand, cortical staining of CLASP1 was more even (Fig. 1L). Finally, we examined the localization of ninein by expansion microscopy, since enrichment of ninein at the cell cortex has previously been described (Lechler & Fuchs, 2007; Lecland et al., 2019). In keratinocytes of the basal layer, concentrated centrosomal staining of ninein was visible (Fig. 1M). In suprabasal layers, centrosomal ninein disappeared, and only cortical ninein was detected. Consistent with earlier studies, ninein was enriched next to desmosomes (Fig. 1M). The cortical enrichment of ninein was not very pronounced in the first two suprabasal layers, but more visible above the third suprabasal layer. Immunofluorescence of microtubules showed ninein present at a subset of microtubule ends, but also near the lateral surface of microtubules (Fig. 1N).

### Microtubule growth towards the cortex in keratinocyte cultures

Since we observed microtubules forming lateral contacts with the cell cortex, as well as end-on contacts of apparently mixed polarity, we investigated directly the growth of microtubule plus-ends in EB3-GFP-transfected keratinocyte cultures, by time-lapse microscopy. Cells were fixed after recording, and stained for immunofluorescence of gamma-catenin, to mark the cortex of the transfected MPEK cells (Fig. 2A, B). By time-lapse microscopy, we observed plus-ends growing from the cytoplasm towards the cell cortex (Fig. 2C), plus-ends growing from the cortex towards the cell center (Fig. 2D), and plus-ends growing in close contact to the cell periphery (Fig. 2E). An average of 35 EB3 comets per cell were tracked near the cortex and each was assigned to one of these three categories. We found that the vast majority of EB3 comets (80%, n=6 cells) were growing from the cytoplasmic center of the cell towards the cortex, whereas only 10% grew in the opposite direction, or closely along the cellular cortex (10%; Fig. 2F).

**Figure 2:**
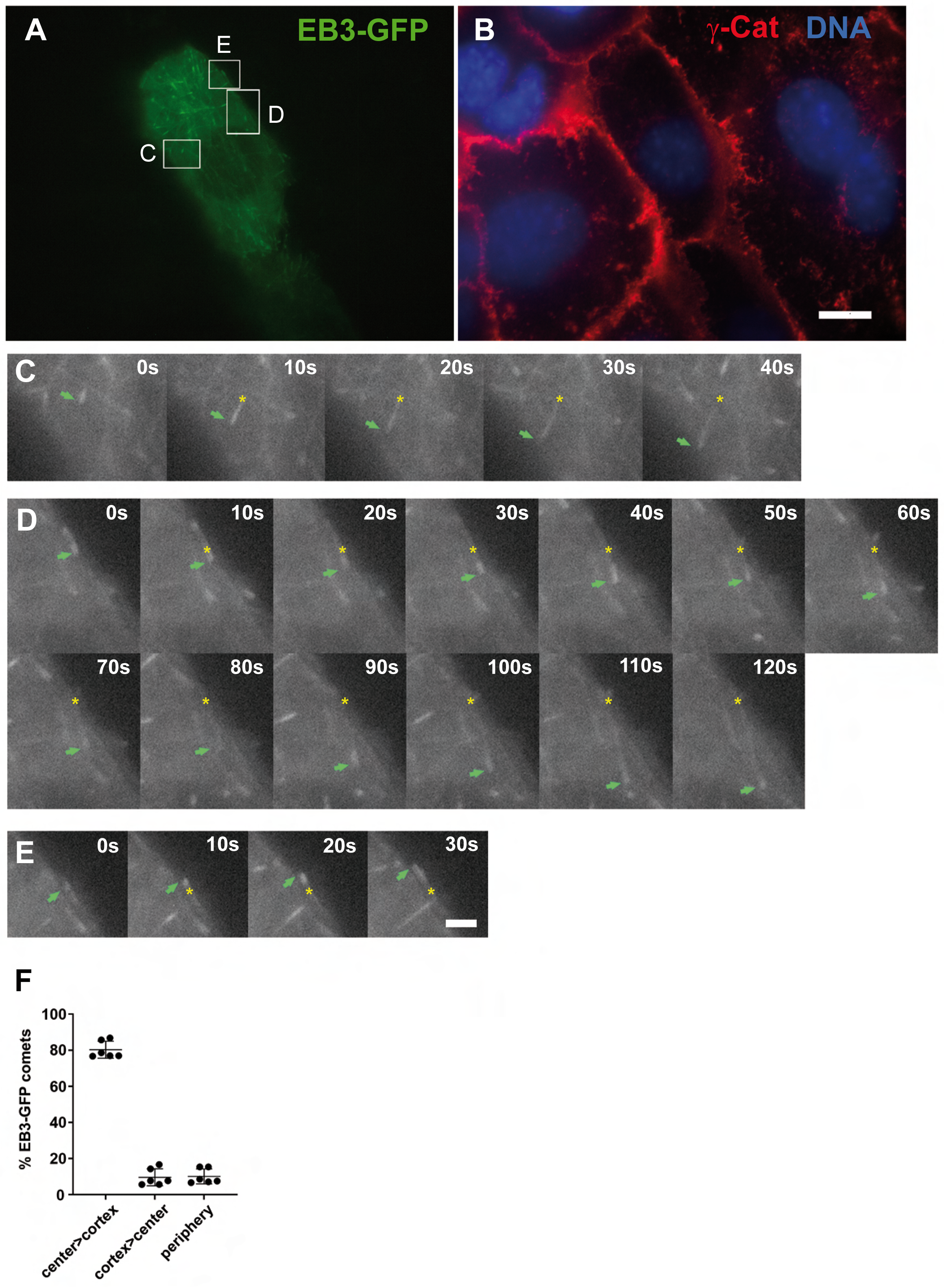
The majority of microtubules grow from the cell center towards the periphery in differentiated MPEK cells. (A) MPEK cell transfected with EB3-GFP, to follow growing microtubule plus-ends. Rectangular frames indicate the areas of time-lapse observations of EB3 comets in (C), (D), and (E). (B) The same cell as in (A), fixed after the time-lapse recording and stained with antibody against gamma-catenin (g-Cat). (C-E) Different areas of the cell in (A), showing EB3 comets in time-lapse recordings at 10s intervals. The growing microtubule-plus ends are indicated by green arrows, the initial starting positions are indicated by yellow asterisks. (F) Percentage of EB3-GFP comets that grow from the cell center towards the cortex (center>cortex), from the cortex towards the cell center (cortex>center), and comets growing along the cell periphery (periphery). Data were obtained from 6 cells. An average of 35 EB3 comets were tracked in each cell near the cortex, and each comet was assigned to one of the three categories. Each data point indicates the percentage of comets in one cell, of one specific category. Bars, (B) 10mm, (E) 2mm.

### Centrosomes are the predominant sites of microtubule re-growth after depolymerization

Because EB proteins localize to growing plus-ends of newly nucleated microtubules and also of fully assembled microtubules that undergo further elongation, the data on the direction of growth of EB3 comets (Fig. 2) don’t allow conclusions on the original sites of microtubule nucleation. We designed therefore an assay of microtubule depolymerization on ice, in the presence of nocodazole, followed by re-nucleation in the absence of the drug, at 37°C. The advantage of this assay is that it reveals, in a sensitive manner, sites where microtubules are nucleated transiently, from a very concentrated soluble pool of depolymerized tubulin, even though these nucleation sites might get lost with time, when the cell re-establishes a balance of polymerized versus soluble tubulin. We examined the dynamic localization of microtubules and microtubule-end markers (Fig. 3A-C), as well as proteins of the microtubule-organizing complex of ninein, dynein, and dynactin in MPEK cells (Fig. 4A, B). The cellular cortex was marked with antibody against desmoplakin (Fig. 3A, Fig. 4B). Before depolymerization (Figs. 3 & 4, control), we detected a subset of microtubules at the cell cortex that largely disappeared after treatment with nocodazole in the cold (0°C, Fig. 3B, Fig. 4A). Nevertheless, a large number of stable microtubules proved to be resistant to this treatment (0°C, Fig. 3B, Fig. 4A). Re-warming the cells in drug-free medium led to the immediate formation of small, but dense centrosomal microtubule asters (2 min, arrows in Fig. 3B, Fig. 4A) that disappeared after prolonged re-warming (8 and 30 min, Fig. 3B, Fig. 4A). Cortical microtubules re-formed upon re-warming (2 min, Fig. 3B), and remained over time (30 min, Fig. 4A). During the process of depolymerization and re-polymerization, the microtubule-nucleating protein gamma-tubulin remained concentrated at the centrosome, while cortical gamma-tubulin was not detectable (Fig. 3A). This was in contrast to gamma-tubulin immunofluorescence in epidermal tissue, where gamma-tubulin disappeared from the centrosome in differentiated keratinocytes of the spinous layer, suggesting that MPEK cultures mimic an incomplete differentiation program of keratinocytes *in vivo*. The microtubule minus-end-stabilizing protein CAMSAP2 concentrated apparently at or near the centrosomes early upon re-warming of MPEK cells (2 min, Fig. 3B, C), but disappeared gradually from these sites at later time points (Fig. 3B, 8-30 min). Meanwhile, cortical enrichment of CAMSAP2 became detectable (Fig. 3B, 8-30 min). Likewise, the marker protein of growing microtubule plus-ends, EB1, was seen highly enriched at the centrosome at 2 minutes, but this accumulation was lost at later time points (Fig. 3C). Altogether, our data suggest early growth of microtubules from the centrosome, but no centrosomal microtubule growth at later time points (8-30 min) or under steady-state conditions (control), consistent with published literature (Lechler & Fuchs, 2007).

**Figure 3:**
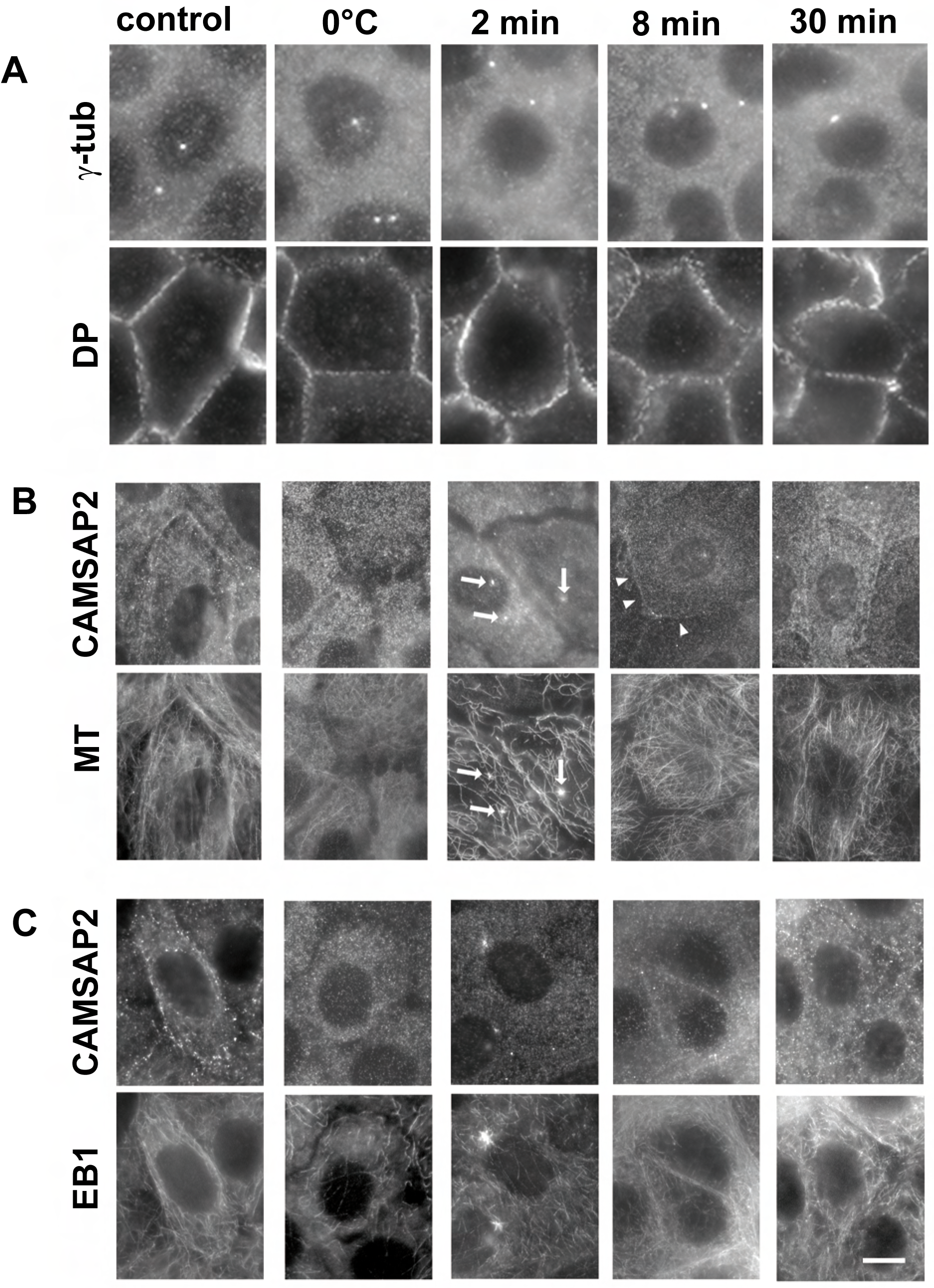
Localization of minus-end and plus-end markers in keratinocyte cultures after microtubule depolymerization and regrowth. MPEK cells were differentiated in culture and stained for immunofluorescence of desmoplakin (DP), alpha-tubulin (MT) and various markers of the microtubule plus-and minus-ends. “Control” indicates cells without any treatment. For the experiment, cells were treated with 3.3 mM nocodazole on ice, followed by removal of the drug on ice (0°C), and subsequent re-warming at 37°C for 2 minutes (2min), 8 minutes (8 min), and 30 minutes (30 min). Localization of (A) gamma-tubulin (g-tub) and desmoplakin (DP), (B) CAMSAP2 and microtubules (MT), (C) CAMSAP2 and EB1. Arrows in (B) indicate the positions of centrosomal microtubule asters, arrowheads indicate the localization of cortical CAMSAP2. Bar, (C) 10mm.

**Figure 4:**
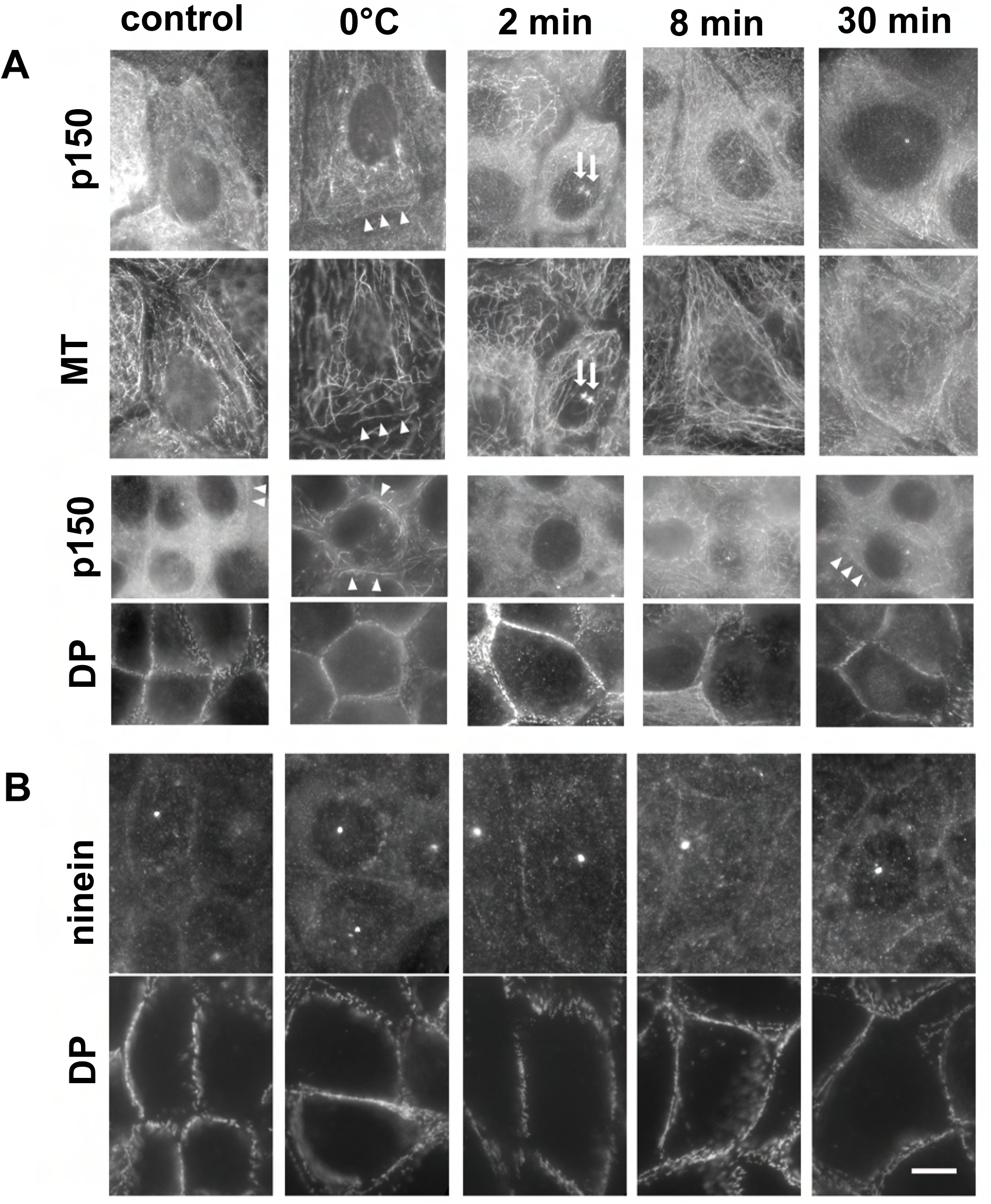
Localization of ninein and its interactor dynactin in keratinocyte cultures after microtubule depolymerization and regrowth. Experimental conditions as in Figure 3. MPEK cells were stained for (A) dynactin subunit p150, microtubules (MT), desmoplakin (DP), and (B) ninein and desmoplakin (DP). Arrowheads in (A) “control” and at 30 min indicate sites of cortical dynactin accumulation, arrowheads at 0°C indicate the co-localization of dynactin with drug-and cold-resistant microtubules at the cortex, and arrows at 2 min indicate the positions of centrosomal microtubule asters. Bar, (B) 10mm.

The microtubule-organizing protein ninein displayed localization at both the centrosome and the cellular cortex, throughout all phases of our experiment, indicating microtubule-independent targeting to both sites (Fig. 4B). As seen for gamma-tubulin (Fig. 3A), centrosomal staining of ninein in differentiated MPEK cells differed from the ninein localization pattern in suprabasal layers of epidermal tissue (Fig. 1M), where ninein was absent from the centrosome. Moreover, p150, an essential subunit of the dynein/dynactin complex, was visible at both the centrosome and the cell cortex, with a peak of increased centrosomal localization early upon microtubule re-growth (2 min, Fig. 4A, arrows). Centrosomal and cortical localization of p150 are consistent with earlier data (Gill et al., 1991; Lecland et al., 2019). A subset of p150 concentrated on drug-resistant, stable microtubules in the cold (Fig. 4A, 0°C, arrowheads). Together with our data on microtubule-end markers (Fig. 3), our experiments on ninein and p150 failed to provide evidence of microtubule nucleation at the keratinocyte cortex, but suggest that the ninein-dependent machinery for cortical microtubule anchorage is permanently present in the differentiated cells, and that CAMSAP2-positive, non-centrosomal microtubules accumulate at the cortex.

### Ectopic expression of ninein at the cell cortex leads to accumulation of known ninein-interactors

Because it has previously been shown that ninein supports anchorage of microtubules, to centrosomal sites in non-differentiated cells, as well as to cortical sites in differentiated keratinocytes (Dammermann & Merdes, 2002; Lecland et al., 2019), we tested directly whether ninein accumulation at the cortex is sufficient to organize a cortical microtubule network, in a cell line that does not organize cortical microtubules under regular conditions. We chose 3T3 mouse fibroblasts, transfected with GFP-ninein that was modified at the carboxy-terminus, by fusing a CAAX sequence motif from KRas. CAAX sequence motifs mediate posttranslational prenylation and anchorage of the protein to the plasma membrane or to the nuclear membrane (Gao et al., 2009). In our experiments, GFP-ninein-CAAX formed large stretches at the plasma membrane. This overexpression of ectopic ninein had no general effect on the centrosome, as marker proteins such as pericentrin or Cdk5rap2 maintained their typical centrosomal localization (Fig. 5A, B). At the cortical sites of GFP-ninein, several previously characterized interactors of ninein accumulated, including Cep170, as well as components and regulators of the dynein/dynactin complex, such as dynein intermediate chain, Lis1, and p150/dynactin (Fig. 5C, H, I, J). Gamma-tubulin, however, failed to accumulate at these cortical sites (Fig. 5D). Because the gamma-tubulin-interaction domain was mapped to amino acids 1-246 in ninein (Delgehyr et al., 2005), we reasoned that the amino-terminal GFP tag might potentially interfere with the ninein-gamma-tubulin interaction. To test this idea, we cloned a construct of ninein with a carboxy-terminal GFP-tag, and we modified an anti-GFP nanobody (Caussinus et al., 2011) with a carboxy-terminal CAAX-tag. Co-transfection of both plasmids yielded cortical enrichment of ninein-GFP, similar to the GFP-ninein-CAAX construct. Under these conditions, we detected cortical enrichment of gamma-tubulin (Fig. 5E). We quantified the gamma-tubulin immunofluorescence intensity within zones of approximately 2mm thickness, along the sites of cortical ninein-GFP, and we detected a significant increase of signal, compared to equivalent zones in control cells, transfected with GFP-CAAX (Fig. 5G). Despite the enrichment of gamma-tubulin, we observed no particular enrichment of microtubules at these cortical sites (Fig. 5F). We noticed, however, that the centrosomal anchorage of microtubules was largely lost, and we further noticed that the percentage of cells displaying EB1 comets emanating from the centrosomal area was significantly reduced (Fig. 5K, L, arrows indicate the origins of centrosomal EB1 comets in non-transfected control cells), compared to control cells expressing GFP-CAAX (90% of control cells with centrosomal EB1 comets, n=260, versus only 36% of cells expressing GFP-ninein-CAAX, n=241). Since the organization of the Golgi apparatus depends on microtubules and dynein/dynactin (Burkhardt et al., 1997; Palmer et al., 2009), we also examined the localization of the Golgi marker protein GM130 in cells expressing cortical ninein. As shown in Figure 5M, this treatment dispersed the Golgi stacks, probably due to sequestration of dynein/dynactin to sites of overexpressed cortical ninein, and partly due to the loss of a focused microtubule network.

**Figure 5:**
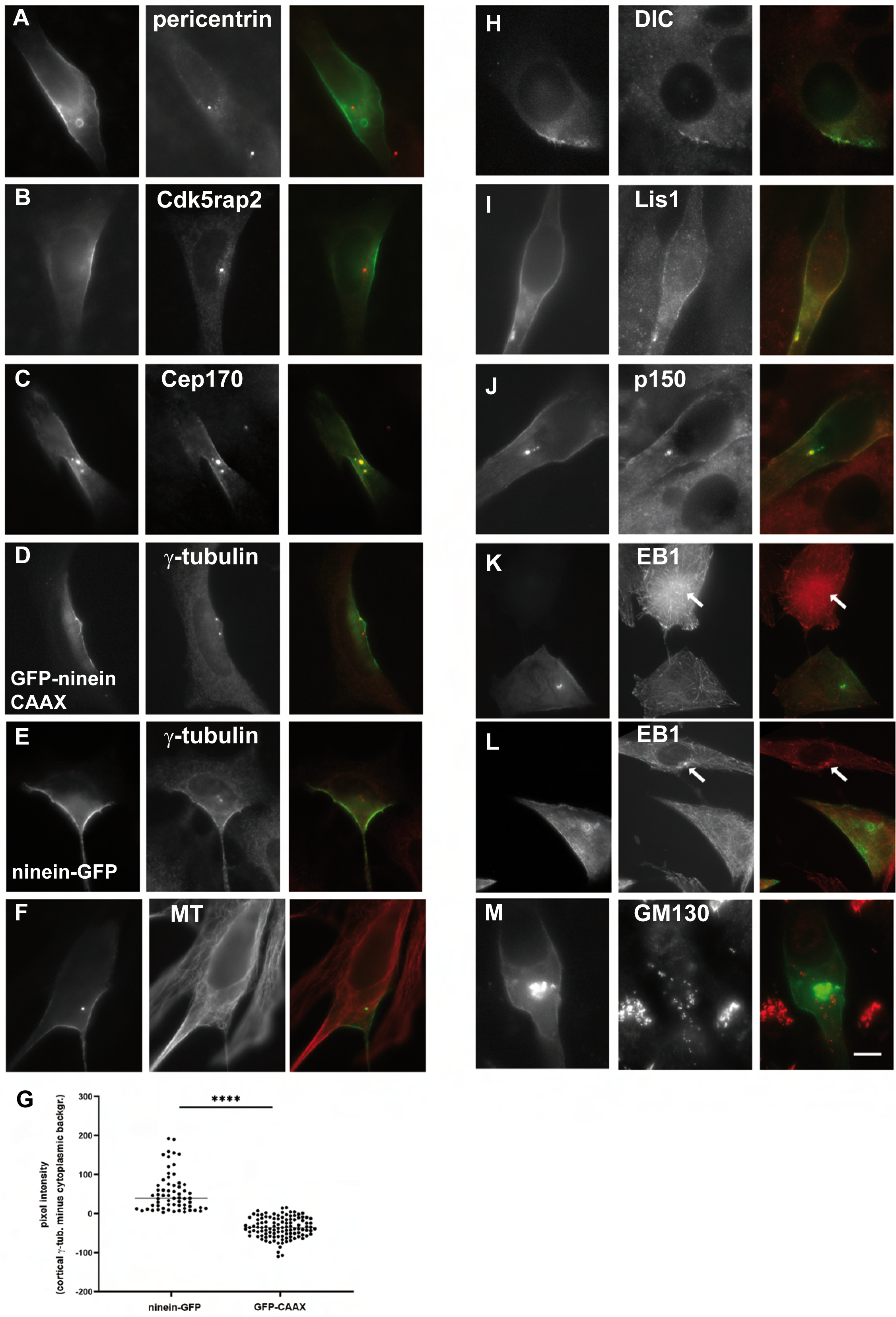
Ectopic ninein at the cell cortex of transfected fibroblasts attracts known ninein-interactors and interferes with centrosomal microtubule organization. 3T3 fibroblasts were transfected with EGFP-ninein-CAAX (A-M, except C, E, F, G), or with anti-GFP nanobody-CAAX in combination with ninein-EGFP (C, E, F, G). GFP fluorescence of tagged ninein is indicated on the left image of each panel. Images in the middle show the localization of (A) pericentrin, (B) Cdk5rap2, (C) Cep170, (D, E) gamma-tubulin, (F) microtubules (MT), (H) dynein intermediate chain (DIC), (I) Lis1, (J) dynactin subunit p150, (K, L) EB1, (M) Golgi marker protein GM130. Images on the right show merged views, with GFP in green and all other markers in red. Arrows in (K, L) indicate radial, centrosomal arrays of EB1 comets. (G) Mean pixel intensities of gamma-tubulin immunofluorescence at a zone of 2mm thickness at the cell cortex, in zones of ninein-EGFP localization (left), or in zones of GFP-CAAX localization, which was transfected as a control. The mean value of cytoplasmic background immunofluorescence of gamma-tubulin was subtracted for each data point, resulting in negative values for part of the GFP-CAAX-transfected cells, since the cortical gamma-tubulin signal was occasionally weaker than the cytoplasmic background. N=63 cells transfected with anti-GFP nanobody + ninein-EGFP, n=111 cells transfected with control GFP-CAAX. P<0.0001. Bar, (M) 10mm.

### Ectopic ninein supports microtubule nucleation and accumulation of severed microtubules

Because cortical ninein-GFP caused accumulation of gamma-tubulin, but no enrichment of cortical microtubules, we tested whether sites of cortical ninein were able to support microtubule nucleation, at least transiently, after cold-induced depolymerization. After 2 minutes recovery from the cold, we detected small stubs of microtubules at cortical sites of ninein-induced gamma-tubulin accumulation, whereas control cells with cortical GFP-CAAX failed to enrich gamma-tubulin and microtubules (Fig. 6A, B). It remains ambiguous whether these cortical microtubules persist, since after re-establishment of steady-state conditions, single microtubules parallel to the cortex could occasionally be detected, but no particular concentration of cortical microtubules with end-on contacts were visible (Fig. 6C).

**Figure 6:**
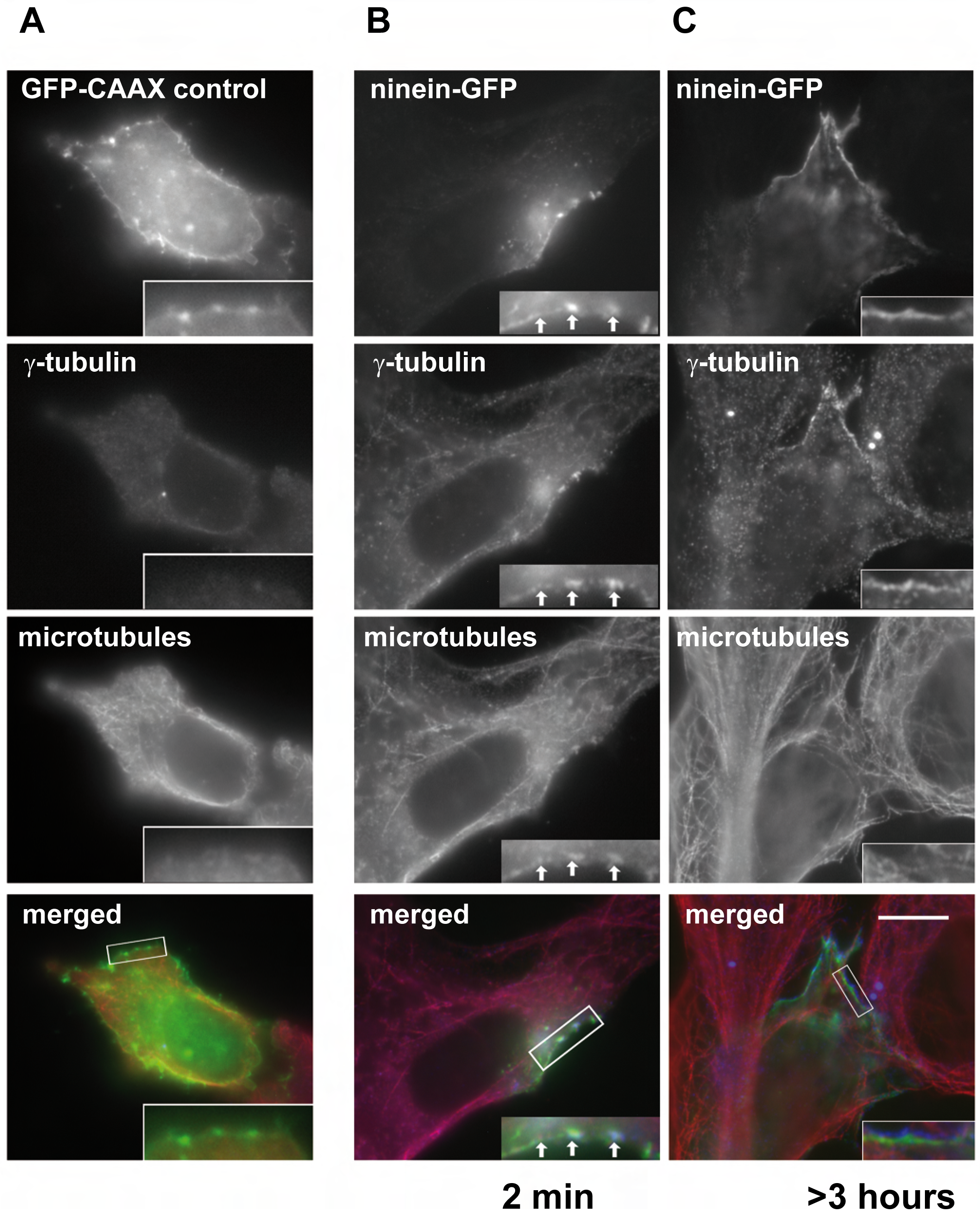
Transient assembly of microtubules at sites of ectopic cortical ninein and gamma-tubulin. (A) 3T3 cell transfected with GFP-CAAX, stained for immunofluorescence of gamma-tubulin and microtubules. Insets show enlarged views of a cortical area, enriched in GFP-CAAX. (B, C) Cells transfected with anti-GFP nanobody and ninein-EGFP, stained as in (A). The cell in (B) was subjected to cold treatment (1h on ice), followed by microtubule re-growth for 2 minutes at 37°C. The cell in (C) was allowed to grow at 37°C for several hours. Insets in (B) show an enlarged area of ninein-EGFP enrichment at the cortex, co-localizing with gamma-tubulin and small microtubule stubs (positions indicated by arrows). (A-C) were stained with monobodies against alpha-and beta-tubulin from guinea pig in combination with goat anti-guinea pig/Alexa 555, and mouse anti-gamma-tubulin in combination with goat anti-mouse/Alexa 647. Merged views of all cells show GFP signal in green, microtubules in red, and gamma-tubulin in blue. Bar, (C, bottom) 10mm.

Because ninein is anchored in a stable manner at the desmosomes of keratinocytes (Lechler & Fuchs, 2007), and because ninein has been characterized as an interactor of the dynein/dynactin complex (Casenghi et al., 2005; Redwine et al., 2017), we considered that cortical microtubules might accumulate by translocation, powered by dynein/dynactin that is immobilized by ninein at the desmosomes. To test this idea, we mimicked experimental conditions in 3T3 cells by immobilizing GFP-ninein-CAAX at the plasma membrane, and by mobilizing microtubules with the help of the severing enzyme spastin. We investigated cells 6 hours after co-transfection with EYFP-spastin, when the microtubule density was diminished due to severing, but with a large number of microtubules still intact. We noticed that spastin-expression yielded cortical microtubule enrichment in cells (Fig. 7A). The degree of enrichment was calculated by determining the ratio of cortical to cytoplasmic microtubule fluorescence. This ratio was significantly higher in cells expressing active spastin, compared to cells expressing an inactive form, EYFP-spastin C530 mut1A (Fig. 7A-C; Lacroix et al., 2010).

**Figure 7:**
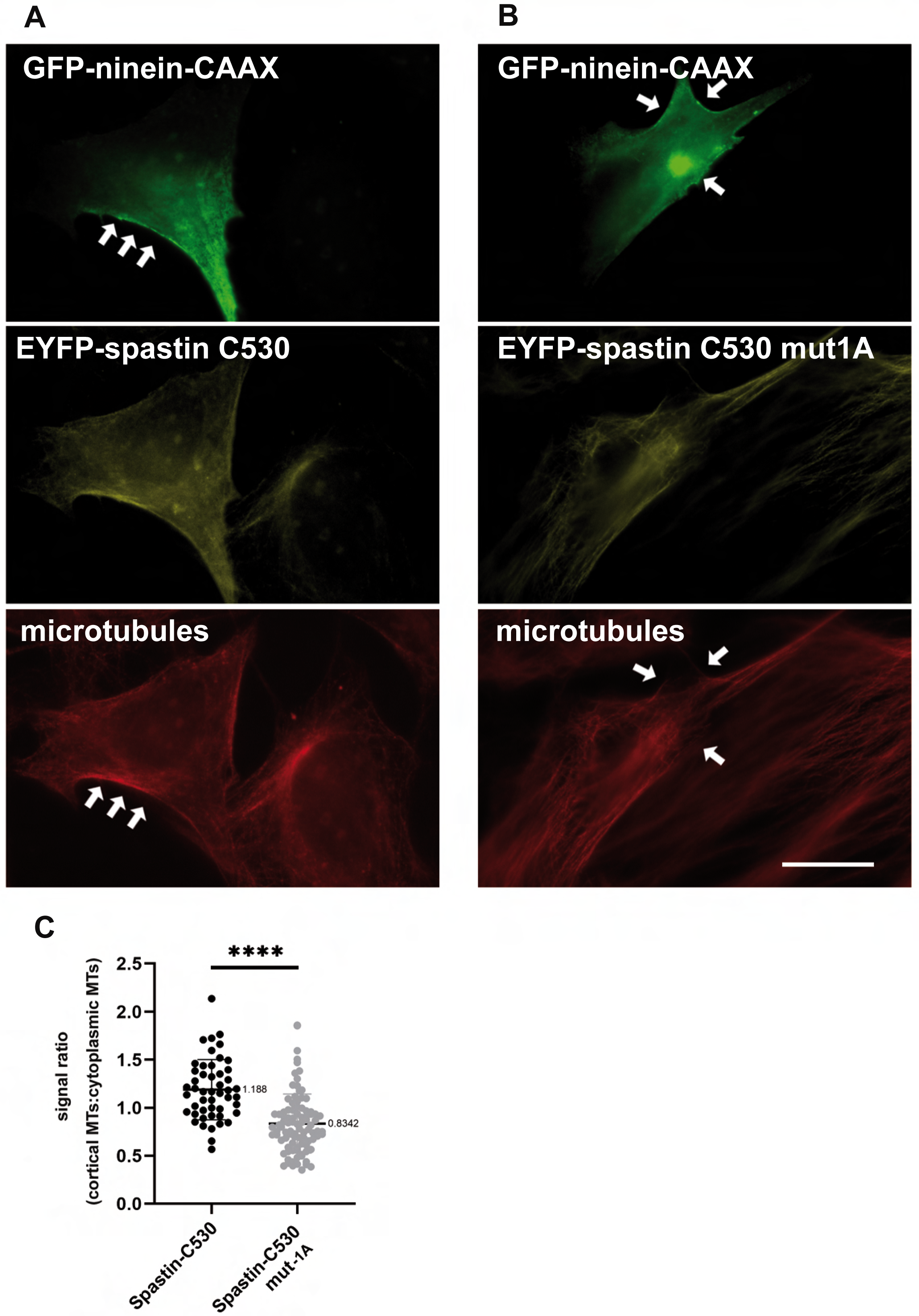
Severing of microtubules by spastin induces the accumulation of microtubules at cortical sites of ectopic ninein. (A) 3T3 cells were co-transfected with EGFP-ninein-CAAX (green) and EYFP-spastin C530 (yellow). After 6h, cells were stained for immunofluorescence of alpha-tubulin (microtubules, red). The EYFP signal was viewed with a 510 nm excitation filter, a Triband dichroic mirror, and a 542 nm emission filter, showing some bleed-through signal from the red channel. Arrows show cortical sites of ectopic ninein and microtubule accumulation. (B) Cells treated as in (A), but using a construct of an inactive form of spastin, EYFP-spastin C530 mut1A. Arrows indicate the sites of EGFP-ninein-CAAX accumulation, largely absent for microtubule accumulation. (C) The ratio of cortical microtubule immunofluorescence to the mean cytoplasmic microtubule immunofluorescence intensity was calculated from cells transfected with spastin C530 (n=49) and inactive spastin C530 mut1A (n=93). P<0.0001. Bar, (B, bottom) 10mm.

## Discussion

In many differentiating cell types, the microtubule network becomes reorganized, with the initial centrosomal focus of microtubules disappearing, followed by anchorage of microtubules at non-centrosomal sites (Dyachuk et al., 2016). In epidermal keratinocytes, non-centrosomal microtubules are anchored at the cellular cortex by desmosomes, initially described by Lechler & Fuchs (2007). In this early study, microtubules were visualized indirectly, by expression of a GFP-tagged microtubule-binding fragment of the protein ensconsin (GFP-EMTB), which raises the question whether ensconsin expression doesn’t favor artificially the formation of cortical microtubules. To circumvent this potential problem, we stained microtubules in the epidermis directly, using antibodies against alpha-tubulin and employing expansion microscopy. Our experimental set-up allowed us to visualize for the first time the contact sites of microtubules at the desmosomes. Our data show that a subset of microtubules in skin keratinocytes is anchored at or near desmosomes, while large numbers of microtubules extend across the cytoplasm in apico-basal or diagonal orientations. This contrasts the findings by Lechler & Fuchs (2007), who showed almost exclusive accumulation of microtubules at the cortex, and only a very low density of microtubules in the cytoplasm. We believe that the previously reported cortical microtubule accumulation can be explained by the expression of GFP-EMTB in the epidermis (Lechler & Fuchs, 2007). Although this ensconsin fragment has been found to bind to microtubules without altering their dynamics (Faire et al., 1999), it is possible that overexpression of GFP-EMTB under the control of the strong keratin 14 promoter may cause artifacts in keratinocytes, since similar overexpression studies in other cell types have shown to stabilize microtubules and to induce the formation of microtubule bundles (Faire et al., 1999).

In our experiments, we detected cortical microtubules whose ends were in contact with desmosomal plaques, besides an equal number of microtubules that ran parallel along the cellular cortex. While these parallel microtubules often localized at around 200 nm distance from desmosomes, it is nevertheless possible that they established molecular contacts with the desmosomal plaques, since the mediator of desmosomal microtubule anchorage, ninein (Lechler & Fuchs, 2007; Lecland et al., 2019), possesses a large rod domain, forming a non-continuous coiled-coil of approximately 1700 amino acids, equaling a predicted length of more than 250 nm. Moreover, ninein interacts with the dynein/dynactin complex at the keratinocyte cortex (Lecland et al., 2019), prolonging the distance for potential microtubule anchorage even further. Potential false-positive co-localization due to misalignment of microscope filter cubes or due to chromatic aberration can be excluded, since maximal deviations of 0.03 mm in X, Y, and 0.1 mm in Z were calculated for expansion microscopy. In addition, we performed immuno-electron microscopy of cortical microtubules in keratinocytes that confirmed our data from expansion microscopy.

Because we detected microtubules with end-on contacts at desmosomes, we wanted to determine the polarity of these microtubules. Immunofluorescence staining of epidermal tissue revealed the presence of microtubule minus-end but also plus-end markers at the cell cortex. We attempted therefore to record microtubule growth in keratinocyte cultures, by following EB3-GFP comets in time-lapse experiments. The low number of microtubules growing from the cellular cortex towards the cell center suggested that microtubule nucleation at the cortex is rare. Consistently, we failed to see proteins of the gamma-tubulin ring complex at the keratinocyte cortex, although the quantity and density of any gamma-tubulin complexes might have been too low to be detected.

When we performed microtubule depolymerization experiments in keratinocytes, followed by regrowth, we noticed that cortical microtubules reformed rapidly within 2 minutes, although EB1 comets were absent from the cortex at this stage, suggesting that these microtubules did not form from cortical nucleation. At the same time, the centrosome was the predominant site of microtubule nucleation, as evidenced by the formation of dense microtubule asters and by the massive accumulation of EB1. Consistent with previously published findings (Lechler & Fuchs, 2007), these asters disappeared within a few minutes, and only non-centrosomal microtubule organization was visible at steady-state conditions. The absence of microtubule nucleation at the keratinocyte cortex was unexpected at first glance, since cortical ninein was permanently present, even in the absence of microtubules (Fig. 4, as well as Lechler & Fuchs, 2007), and since ninein was found to bind gamma-tubulin ring complexes (Delgehyr et al., 2005). To our surprise, we succeeded in redirecting gamma-tubulin to the plasma membrane under specific experimental conditions in non-keratinocyte cells, when we sequestered ninein-GFP to the cortex by a CAAX-tagged nanobody against GFP. Moreover, from these sites containing ninein-GFP and gamma-tubulin, short microtubule stubs formed after cold-induced depolymerization and re-polymerization. However, this microtubule formation occurred only transiently, suggesting that factors that anchor or stabilize microtubules were absent from these sites. In a similar context, ectopic sites of gamma-tubulin were created in keratinocytes, and it was shown that separate proteins are necessary for activation of microtubule nucleation and for microtubule anchorage (Muroyama et al., 2016). In particular, it was demonstrated that microtubule-organizing centers that nucleate new microtubules can only retain them if the anchorage factor NEDD1 is present (Muroyama et al., 2016). It is therefore possible that in our own experiments on ninein-GFP-dependent gamma-tubulin recruitment at the cell cortex, NEDD1 or other factors were not present in sufficient quantities to allow for permanent anchorage of microtubules.

Altogether, our data indicate that cortical microtubules in keratinocytes can be generated by cortical nucleation, but likely only in small amounts and at low stability (Fig. 8A). In an alternative pathway (Fig. 8B), microtubules may be nucleated at non-cortical sites, such as the centrosome, followed by release and translocation towards the cortex (Keating et al., 1997). Alternatively, free microtubules may be created by severing microtubule fragments from longer microtubules, with the help of severing enzymes such as katanin or spastin, followed by translocation. Mechanisms that involve severing and translocation have been suggested to support cortical microtubule organization in *Arabidopsis*, and to facilitate axon outgrowth in neurons (Ahmad et al., 1999; Burk et al., 2001). Translocation has also been proposed as a mechanism for the reorganization of microtubules from the centrosome to the cellular cortex in inner and outer pillar cells of the organ of Corti (Mogensen et al., 2000). In this experimental system, it has been suggested that ninein is released from the centrosome and kept attached to the minus-ends of translocating microtubules during the reorganization process. Different from the proposed mechanism in pillar cells, we favor a model for keratinocytes in which ninein is stably anchored at the cortex by desmosomes. Due to its ability to bind to the dynein/dynactin complex, it can “reel in” free microtubules with the help of the minus-end-directed motor activity of dynein (Fig. 8B). Immobilized by ninein and desmoplakin, dynein would pull on microtubule plus-ends, and translocate the microtubules to various degrees, thereby creating cortically anchored microtubules with various orientations: at the start of the translocation process, the microtubule plus-end would be in contact with ninein/dynein/dynactin at the desmosome, a partially reeled-in microtubule would have a lateral contact with the desmosome, and a fully reeled-in microtubule would have its minus-end in contact with the desmosome (Fig. 8B, bottom, from the right to the left). As part of the process of centrosomal release and translocation, free microtubule minus-ends may be stabilized by CAMSAP2, as seen in our immunofluorescence experiments. A stabilized cortical microtubule array can then facilitate the assembly and maintenance of functional desmosomes and the fusion of lamellar bodies, in keratinocytes of the spinous and granular layers, and thereby contribute to the formation of an intact epidermal barrier (Lecland et al., 2019).

**Figure 8:**
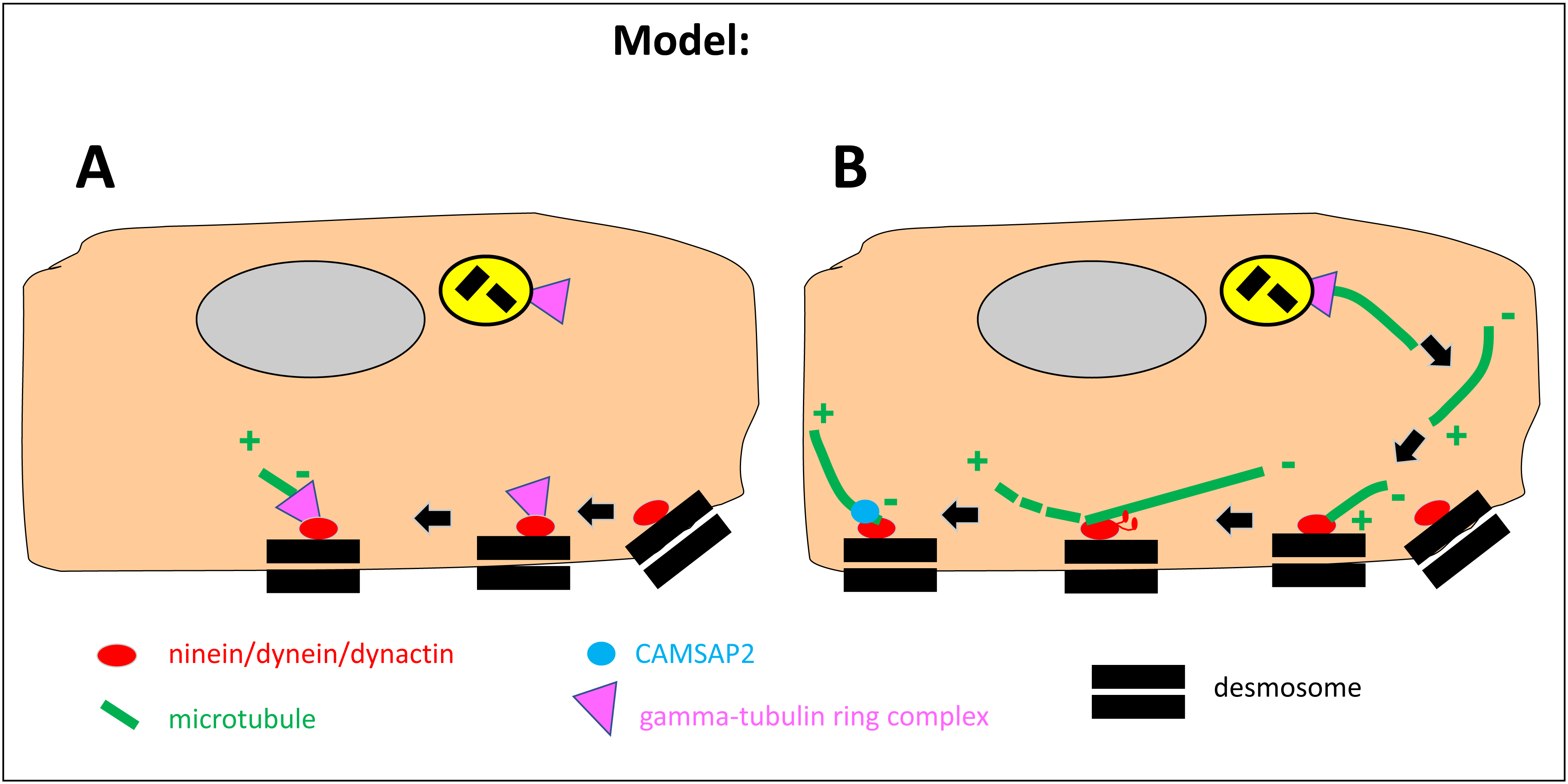
Model of different pathways of cortical microtubule organization in epidermal keratinocytes. (A) Ninein is bound to desmosomes via desmoplakin. Microtubule nucleation takes place at the cortex, from ninein-bound gamma-tubulin ring complexes. (B) Microtubules are nucleated at non-cortical sites in the cell center, such as the centrosome. Microtubules are released and translocated towards the cortex. Translocation at or near the cortex is mediated by a pulling force, created by the minus-end-directed motor activity of the dynein/dynactin complex. Dynein/dynactin itself is anchored and activated by ninein, at the sites of desmosomes. Cortical microtubule minus-ends are stabilized by CAMSAP2. The pathway in (A) creates microtubules with minus-ends anchored at the cortex. The pathway in (B) can create cortical microtubules with plus-end attachment at the desmosome, microtubules that are oriented parallel to the cortex, and microtubules with minus-end attachment at the desmosome (indicated from the right to the left). Both pathways (A and B) may operate in parallel.

## Acknowledgements

The authors would like to thank Drs. L. Haren and C. Bierkamp for critically reading this manuscript. L. Haren, C. Bierkamp, J. Benedetti, T. Gilbert, C. Gorlt, and K. Garreau (University Toulouse III) are thanked for technical help and constructive discussions. C. Jahnke and V. Henriot (Institut Curie, Orsay) are thanked for providing spastin constructs. K.G. was the recipient of a PhD scholarship from the China Scholarship Council (No. 202006220048). This work was further supported by salary support from the Centre National de la Recherche Scientifique, France.

---

**Supplemental Figure 1:**
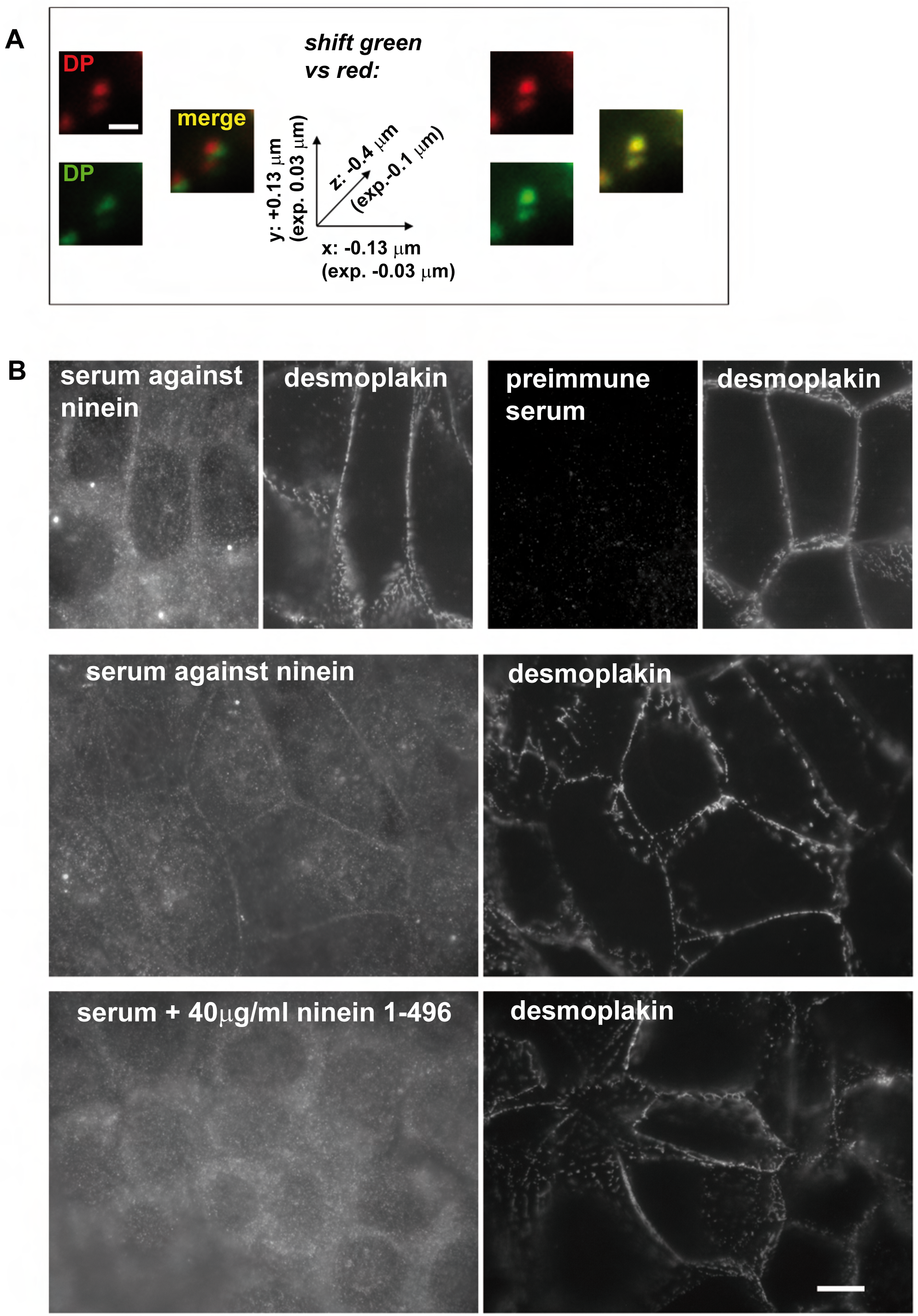
(A) Deviations of filter cube alignments and chromatic aberration lead to lateral and vertical shifts of images recorded in different channels. Desmosomes in MPEK cells, stained with primary antibody against desmoplakin (DP), followed by staining with a combination of secondary antibodies, goat-anti-mouse Alexa 488 and goat-anti-mouse Alexa 568 (left). Z-stacks of images at 0.2mm distance were recorded, and enlarged areas of the desmosomes were subsequently aligned by manually shifting recordings of the individual channels in Adobe Photoshop, until maximal overlap was reached (right). The deviations were calculated in X, Y, and Z, and displayed in the graph (middle). For expanded samples, the deviations can be divided by the expansion factor of 4.1. (B) Characterization of a rabbit antiserum against a hexa-histidine-tagged fusion protein of ninein amino acids 1-496. Top row: differentiated MPEK cells were stained by immunofluorescence using this serum, or the corresponding pre-immune serum from the same rabbit, in combination with mouse-anti-desmoplakin, to stain the centrosome and the cellular cortex. Middle and bottom rows: same staining as in top row images, but in the presence of mounting concentrations of the ninein 1-496 fusion protein (bottom row showing 40mg/ml ninein 1-496), leading to competition with intracellular ninein immunofluorescence. The excess of the fusion protein leads to loss of centrosomal and cortical ninein staining, underlining the specificity of this antiserum. Bars, (A) 1mm, (B) 10mm.

